# Multifunctional semiconducting polymer dots as a miRNA delivery system for osteogenic differentiation and stem cell tracking

**DOI:** 10.1101/2025.08.06.668851

**Authors:** Ping Song, Jørgen Kjems

**Affiliations:** Center for DNA Nanotechnology (CDNA) and Interdisciplinary Nanoscience Center (iNANO), Center for RNA Therapeutics towards Metabolic Diseases (RNA-META), Aarhus University, Gustav Wieds Vej 14, DK-8000 Aarhus C, Denmark; Department of Molecular Biology and Genetics, Aarhus University, 8000 Aarhus C, Denmark

**Author notes:** Correspondence: Jørgen Kjems, Interdisciplinary Nanoscience Center, Department of Molecular Biology and Genetics, Aarhus University, Gustav Wieds Vej 14, DK-8000 Aarhus, Denmark.

## Abstract

Small interfering RNAs (siRNA) and microRNAs (miRNA) are potent gene regulators through RNA interference (RNAi). In this study, we developed a multifunctional theranostic carrier for small RNA delivery and cell tracking using a polymer based nanoparticle composed of the lipid-like material S14, Pluronic F127 and semiconducting polymer MEH-PPV (SPM-dots). The resulting SPM-dots showed efficient siRNA transfection and labeling of human mesenchymal stem cells (hMSCs). Delivery of miR-29b to hMSCs significantly enhanced osteogenic differentiation. Furthermore, hMSCs transfected with SPM-dots displayed strong fluorescent signals in vivo after subcutaneous transplantation. These findings indicate that SPM-dots hold great potential as carrier for gene delivery and cell tracking applications.

## Introduction

Mesenchymal Stem Cells (MSCs) are multipotent pericytes widely used in stem-cell based therapies, including immune modulation and tissue regeneration^1^. Bone marrow-derived MSCs can differentiate into multiple cell types—such as adipocytes, chondrocytes, fibroblasts, and osteoblasts—through mechanisms controlled by different gene regulatory pathways^2,3^. MicroRNAs (miRNAs) have gained recognition for directing stem cell differentiation^4^; Several miRNAs modulate the balance between osteoclastogenesis and osteoblastogenesis and are key regulators of bone regeneration and osteoporosis^5,6^. Among these, miR-29b promotes osteogenesis by repressing osteogenic inhibitors such as β-catenin interacting protein (CTNNBIP1), and transforming growth factor beta 3(TGF-β3), as well as by modulating the accumulation of extracellular matrix protein during the mineralization stage ^7–9^.

Non-viral delivery systems often face challenges in effectively transfecting stem cells, particularly mesenchymal stem cells (MSCs). Consequently, there is a strong need for the development of more efficient delivery systems. Various approaches—including peptides^10,11^, polymers and lipids^12,13^—have been explored for siRNA and miRNA delivery. Recent years, lipid-like materials have been synthesized to improve siRNA delivery, with their efficiency and safety evaluated in mice, rats and nonhuman primates^14^. Several studies have employed lipid-like material based nanoparticles to deliver siRNA in disease models for liver diseases, arthritis and cancer^15–17^. These studies suggest that lipid-like materials with excellent biological properties hold great potential for siRNA/miRNA delivery.

Another challenge in stem cells-based therapy is the development of efficient labeling and tracking techniques that provide detailed information on biodistribution, cell migration and viability. Poly(2-Methoxy, 5-(2’-ethyl-hexyloxy-p-phenylene-vinylene) (MEH-PPV) is a hydrophobic electroluminescent polymer widely used as a semiconductor in light-emitting diodes (OLED), photovoltaic devices and biosensors due to its excellent electrical and spectroscopic properties^18,19^. Through nanotechnology, compounds such as MEH-PPV— previously considered unsuitable for biological applications—can now be adapted for biomedical use. MEH-PPV has attracted significant interests in biological materials science and is increasingly used in semiconducting polymer nanoparticles that possess a π-conjugated structure as excellent fluorescent probes ^20–22^. For example, semiconducting polymer nanoparticles synthesized based on PFPL-(poly(fluorenylene phenylene) and PPE-NH_2_ (amine-containing poly(phenylene ethynylene)) have been employed for plasmid DNA^23^ and siRNA^24,25^ delivery.

In this study, we formulated ultra-bright semiconducting polymer dots, referred to as SPM-dots, through hydrophobic interactions among S14 (a lipid-like material) [S]^17^, Pluronic F127 [P], and the semiconducting polymer MEH-PPV[M]. SPM-dots demonstrated high transfection efficiency in hMSCs, with transfection of SPM-dots/miR-29b significantly enhancing osteogenic differentiation. Furthermore, the optical properties of SPM-dots enabled tracking of transfected cells following subcutaneous implantation in mice.

## Materials and Methods

### Materials

Poly(2-Methoxy, 5-(2 ‘-ethyl-hexyloxy-p-phenylene-vinylene) (MEH-PPV), spermidine, Pluronic F127 and 1,2-epoxytetradecane (ETD) were purchased from Sigma-Aldrich. sequence of the miRNA-29b (hsa-miR-29b) was acquired from miRBase Sequence Database, with sense strand: 5′-UAGCACCAUUUGAAAUCAGUGUU-3′; siRNA against GFP (siGFP) was purchased from Ribotask (Denmark), sense 5′-GACGUAAACGGCCACAAGUTC-3′, and antisense 5′-ACUUGUGG CCGUUUACGUCGC-3′; The negative control siRNA (siNC) was provided by Genepharm, Shanghai, China, sense, 5′-UUCUCCGAACGUGUCACGUTT-3′, and antisense, 5′-ACGUGACACGUUCGGAGAATT-3′. Cy3 labeled siGFP was used for cell uptake studies. Metrigel (BD, Bioscience).

### Synthesis of Lipid-like material

The lipid-like material S14 was prepared following established protocols from our laboratory^17^. Specifically, 1,2-Epoxytetradecane (ETD) and spermidine were combined at a molar ratio of 4:1 (ETD:spermidine) and subjected to stirring at 90°C for 48 hours. The crude product was subsequently purified by column chromatography using gradient elution method, transitioning from CH_2_Cl_2_ to a mixture of CH_2_Cl_2_/MeOH/NH_4_OH (75:22:3). The purified product was kept at −20°C before further use.

### Preparation of SPM-dots

The Semiconducting Polymer dots (SPM-dots) were prepared by nanoprecipitation method. In this process, S14 was dissolved in absolute ethanol at the concentration of 100mg/ml,while MEH-PPV and Pluronic F127 were both dissolved in Tetrahydrofuran (THF) at 50mg/ml. 10 μl S14, 80 μl Pluronic F127 and 20 μl MEH-PPV solutions were combined and rapidly injected into acetate buffer (200mM, pH=5.4) under continuous stirring, facilitating the assembly of polymer dots spontaneously. The polymer dots solution was subjected to dialysis using a 1kDa molecular weight cut-off membrane against the same buffer. To encapsulate siRNA or miRNA, 20 μM of siRNA or miRNA solution was mixed with empty SPM-dots at a weight ratio of 7.5:1 (S14: siRNA/miRNA) and the mixture was incubated at room temperature for 15 min.

### Characterization of the SPM-dots

The hydrodynamic diameter and zeta potential of the SPM-dots were assessed at 25°C using dynamic light scattering (DLS) and laser Doppler electrophoresis on a Zetasizer Nano ZS at 25°C (Malvern Instruments, Malvern, UK). To visualize morphology, Transmission electron microscopy (TEM) was performed. The grid was prepared by adding 5 μL of particle suspension to a carbon-coated 200 mesh copper grid (Microscopy products for Science and Industry, USA) for 1 min. Excess fluid was removed with filter paper, and the grid was immediately stained using uranyl formate.

The siRNA/miRNA encapsulation efficiency was determined with RiboGreen reagent (Invitrogen, Copenhagen) as per the supplier’s protocol. Equal volumes (50 μL) of SPM-dots/siRNA complexes and RiboGreen working solution (1:100 in TE buffer) were mixed and incubated for 5 min. Fluorescence was then quantified using a FLUOstar OPTIMA plate reader (BMG, Labtechnologies) with excitation/emission wavelengths set at 480 /520 nm, respectively. The fluorescence emission profile of the SPM-dots was obtained using a Horiba Jobin Yvon Fluoromax-3 fluorimeter, with excitation at various wavelengths. Absorption spectra of the polymer dots in ultrapure water was measured by a NanoDrop ND-1000 spectrophotometer (Nanodrop, Wilmington, DE, USA).

### Cell culture and Osteoblastic Differentiation

Primary bone marrow-derived primary hMSCs were obtained from ATCC (Manassas, VA, USA), while GFP-expressing transgenic hMSCs were a generous gift Dr. Moustapha Kassem (University of South Denmark, Odense, Denmark). Both cell types were cultured in Minimum Essential Medium (MEM) supplemented with 10% fetal bovine serum and 1% penicillin-streptomycin, maintained at 37°C in 5% CO_2_ and 100% humidity. To induce osteogenesis, cells were exposed to induction medium containing 10 mM β-glycerophosphate, 0.2 mM l-ascorbic acid and 10^−7^ M dexamethasone.

### Cellular uptake studies

hMSC cells were seeded into 8-well chambers (SARSTED, Germany) at a density of 5×10^4^ cells/well and allowed to adhere overnight. The next day, the cells were incubated with SPM-dots/Cy5-labeled siRNA (Cy5-siRNA) at a final concentration of 50 nM siRNA for 4 hours at 37 °C. The cells were fixed for 15min with 4% paraformaldehyde at room temperature, rinsed three times with PBS, and stained 15 min at 37 °C with Alexa Fluor® 488-WGA for membrane labeling and hochest 33342 (Molecular Probes) for nucleus staining. Fluorescent imaging was performed using a Zeiss LSM 710 confocal microscope (Zeiss, Germany). For uptake quantification, hMSCs were seeded at 5×10^4^ cells per well in 24-well plate and cultured overnight. The cells were then treated with SPM-dots/Cy5-labled siRNA (50 nM) for 4 h. The cells were detached using 0.05% Trypsin-EDTA (Gibco, Invitrogen) and examined by Flow cytometry (Becton Dickenson).

### Cell viability

Cell viability was evaluated using the AlamarBlue assay (Molecular Probes, Life Technologies) following the manufacturer’s guidelines. hMSCs were seeded in a 96-well plate at a density of 5×10^3^ cells per well and allowed to adhere overnight. The next day, cells were treated with 100 μL of fresh medium containing SPM-dots/miRNA-NC at the final miRNA concentration of 50 nM. Formulations were tested across different S14: miRNAweight ratios (2.5, 5, 7.5 and 10). After 24-hours incubation, the cells were gently washed with PBS and incubated with 10 % dilution of AlamarBlue reagent in culture medium for 2 h. Fuorescence intensity of the supernatant was recorded using a plate reader (FLUOstar OPTIMA, Moritex BioScience) with excitation at 540 nm and emission at 590 nm.

### GFP gene silencing efficacy of SPM-dots

Transgenic mesenchymal stem cells expressing GFP were seeded at a density of 5×10^4^ cells/well in a 24-well plate. After overnight incubation, cells were treated with SPM-dots/siGFP at a final concentration of 50 nM for siGFP. After incubation overnight, cells were replaced with fresh media and incubated for another 24 h. The GFP silencing was visualized and imaged by microscope (OLYMPUS IX71). To quantify GFP silencing, cells were harvested using 0.05% Trypsin-EDTA and subsequently analyzed by flow cytometry (Becton Dickenson).

### Expression level of osteogenic markers

hMSC cells were plated in 24-well plate (5×10^4^ cells/well) overnight. Then, SPM-dots/miR-29b and SPM-dots/miR-NC were incubated with hMSCs at a final concentration of 50 nM for miRNA. The cells were exposed to induction medium for osteoblastic differentiation for 14 days. To evaluate gene expression, total RNA was isolated with Trizol reagent (Invitrogen), following the manufacturer’s protocol. cDNA was synthesized using Superscript® II Reverse Transcriptase kit (Invitrogen). The resulting cDNA was the template for quantitative PCR, performed with the SYBR® Green kit (Invitrogen) on a LightCycler® 480 Real-Time PCR System (Roche). The quantitative results represent the mean of triplicates per experiment. Primers used for qPCR are listed in Supplementary Material (Table S1).

### Alizarin red Staining

In vitro mineralization was assessed by Alizarin red S staining, as described^26^. Briefly, after 14 days of osteogenic induction, cells were gently rinsed with PBS and fixed in 80% ethanol pre-chilled to −20 °C for 45 min. Following fixation, the cells were rinsed with dH_2_O, and incubated with AR-S solution (20 mg/mL, pH 4.2; Sigma-Aldrich, St. Louis, MO) for 20 minutes at room temperature. To minimize the backgroud staining, the plates were rinsed twice with dH_2_O and washed three times with PBS. For quantitative analysis, the bound AR-S was solubilized in 10% (w/v) cetylpyridinium chloride for 20 min and placed on a shaker (100 rpm) at room temperature. Absorbance was measured at 570 nm using a NanoDrop ND-1000 spectrophotometer (Nanodrop, Wilmington, DE, USA), with each sample measured in triplicate.

### ALP activity

Alkaline phosphatase (ALP) activity was measured by using p-nitrophenyl phosphate (Sigma) as substrate^27^. Briefly, relative cell number was measured by AlamarBlue assay (Molecular Probes, Life Technologies) according to the manufacturer’s protocol on day 7. Cells were then washed with Tris-buffered saline, fixed in 3.7% formaldehyde, 90% ethanol for 30 seconds at room temperature, incubated with substrate (1 mg/ml of p-nitrophenyl phosphate in 50 mM NaHCO_3_, pH 9.6 and 1 mM MgCl_2_) at 37 °C for 20 minutes, and the absorbance was measured at 405 nm, using FLUOstar Omega multimode microplate reader.

### Stem cell labeling and *in vivo* stem cell tracking

For cell labeling, 1×10^6^ hMSC cells were transfected with SPM-dots /miRNA at a final concentration of 50 nM for miRNA and incubated at 37 °C overnight. The cells were harvested and resuspended in PBS (1% Heparin). Different concentrations of cells were mixed with Matrigel and implanted subcutaneously on the back of a Balb/c mouse. Briefly, 1×10^5^, 5×10^4^, 1×10^4^ cells, and 5×10^3^ cells (suspended in Matrigel/PBS=1:1) were injected subcutaneously at the volume of 100 μl. 100 μl. PBS/Matrigel solution was injected as negative control. After 1 h post-injection, the mouse was scanned using an IVIS® 200 imaging system (Xenogen, Caliper Life Sciences, Hopkinton, MA, USA) under 2.5% isoflurane.

### Statistics

Paired Student’s T-test was performed for comparison of two groups for statistics analysis. Significant differences are considered at a level of *P < 0*.*05*.

## Results

### Characterization of the SPM-dots

The SPM-dots were prepared using the ethanol injection method and self-assembled into a core-shell structure via hydrophobic interactions among S14, Pluronic F127 and MEH-PPV. As shown in Figure 1, the hydrophobic, fluorescence-emitting polymer MEH-PPV formed the core of the polymer dots through interactions with the hydrophobic chains of Plutonic F127 and S14. The synthesized SPM-dots exhibited a diameter of 164.7 ± 2.28 nm and a zeta potential of +41.2 ± 4.17 mV, with excellent uniformity indicated by a polydispersity index of 0.14±0.011. Transmission electron microscopy (TEM) analysis confirmed the well-dispersed spherical morphology of the nanoparticles (Figure 1). The particle size observed by TEM of 82 ± 16.1 nm was smaller than the hydrodynamic size measured by DLS, likely due to sample shrinkage during sample preparation.

**Figure 1.**
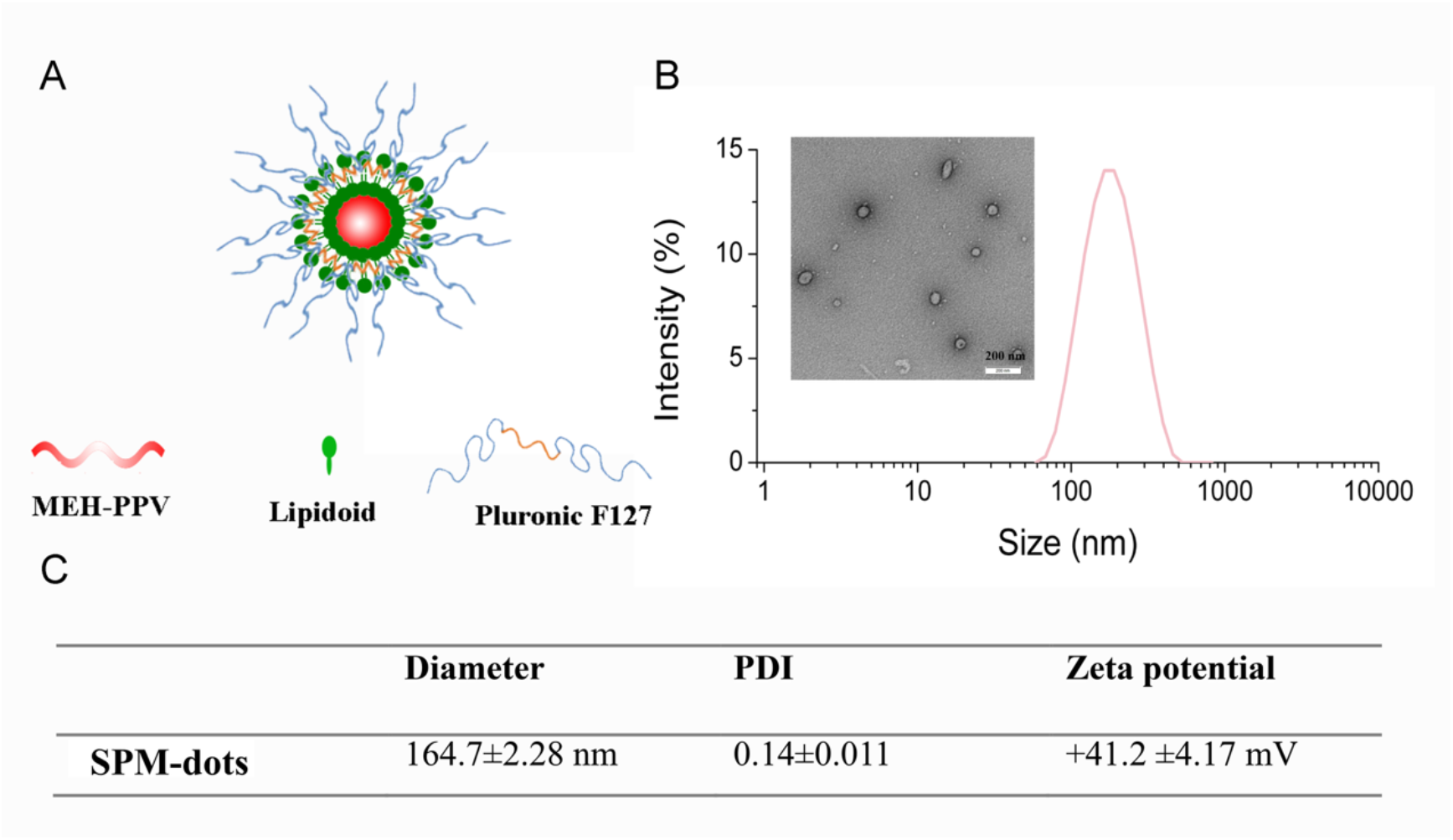
Characterization of the SPM-dots. (A) Illustration of the structure of SPM-dots. (B) Size of SPM-dots measured by TEM and DLS. (C) Size, polydispersity (PDI) and zeta potential of SPM-dots. Data are expressed as mean ± SD (n=3).

The optical properties of SPM-dots were characterized by UV-vis absorbance (Figure 2A) and fluorescence spectra (Figure 2B). The UV-vis spectrum showed a strong absorption peak at approximately 500 nm. Fluorescence emission spectra were recorded at excitation wavelengths ranging from 470 nm to 540 nm, with a prominent emission peak around 590 nm at 500 nm excitation. The fluorescence intensity decreased with increasing excitation wavelength; however, no significant shift in the emission peak was observed across the excitation range.

**Figure 2.**
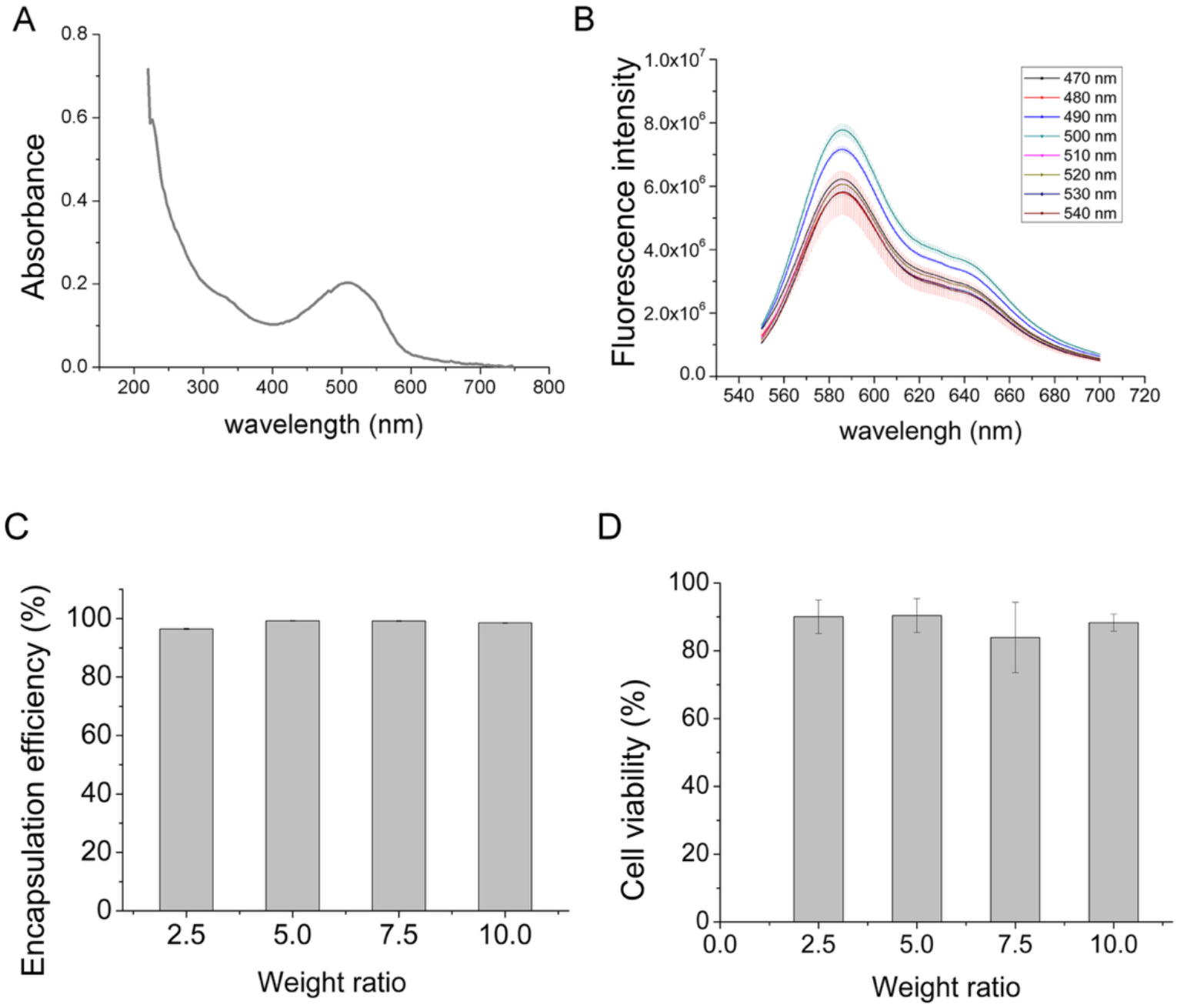
Photophysical behavior, drug encapsulation, and cytocompatibility of SPM-dots(A) UV-vis absorption spectrum of SPM-dots and (B) fluorescence emission spectra of the SPM-dots with excitation wavelength from 470nm to 540nm in 10 nm increments. Each bar represents the mean of fluorescence intensities (n=3) (C) miRNA encapsulation efficiency of SPM-dots at the weight ratio (S14/miRNA, w/w) 2.5 ~10. Data was expressed as mean ± SD (n=3). (D) Cell viability of SPM-dots on hMSCs evaluated by Alarmablue assay after 24 hours incubation at indicated weight ratios. Data was expressed as mean ± SD (n=4).

The miRNA encapsulation capacity of SPM-dots was evaluated at different weight ratios using RiboGreen reagent (Invitrogen, Copenhagen). As shown in Figure 2C, encapsulation efficiency exceeded 90% across all tested ratios (2.5, 5, 7.5 and 10 corresponding to N/P ratios of 2, 4, 6, 8), likely due to the protonatable amine groups of S14. Cytotoxicity of SPM-dots/siRNA complexes was assessed in hMSCs at the same weight ratios after 24 h transfection. As shown in Figure 2D, no significant toxicity was observed at any ratio.

To access the cytotoxicity of SPM-dots, hMSC cells were treated with SPM-dots/miNC (Negative control miRNA) at the weight ratios (S14: miRNA) of 2.5, 5, 7.5, 10 and compared with untreated controls after 24 hours. AlarmarBlue assay results showed approximately 90% cell viability across all ratios, indicating good biocompatibility (Figure 2D). A weight ratio of 7.5 was selected for subsequent experiments, balancing encapsulation efficiency, stability of RNA complexes and low toxicity.

### Cell delivery

To evaluate cellular uptake efficiency, Cy5-labeled siRNA was encapsulated into SPM-dots and transfected into hMSCs, with Lipofectamine2000(Lipofect) as a positive control. Confocal microscopy revealed efficient internalization of Cy5-siRNA delivered by SPM-dots after 4 hours of transfection. Notably, higher Cy5-siRNA fluorescence intensity was observed SPM-dots-treated cells compared to Lipofect-transfected cells (Figure 3A). Flow cytometry analysis confirmed these findings, showing greater accumulation of Cy5-labeled siRNA in SPM-dots-transfected cells (Figure 3B, C), demonstrating superior transfection efficiency relative to Lipofectamine.

**Figure 3.**
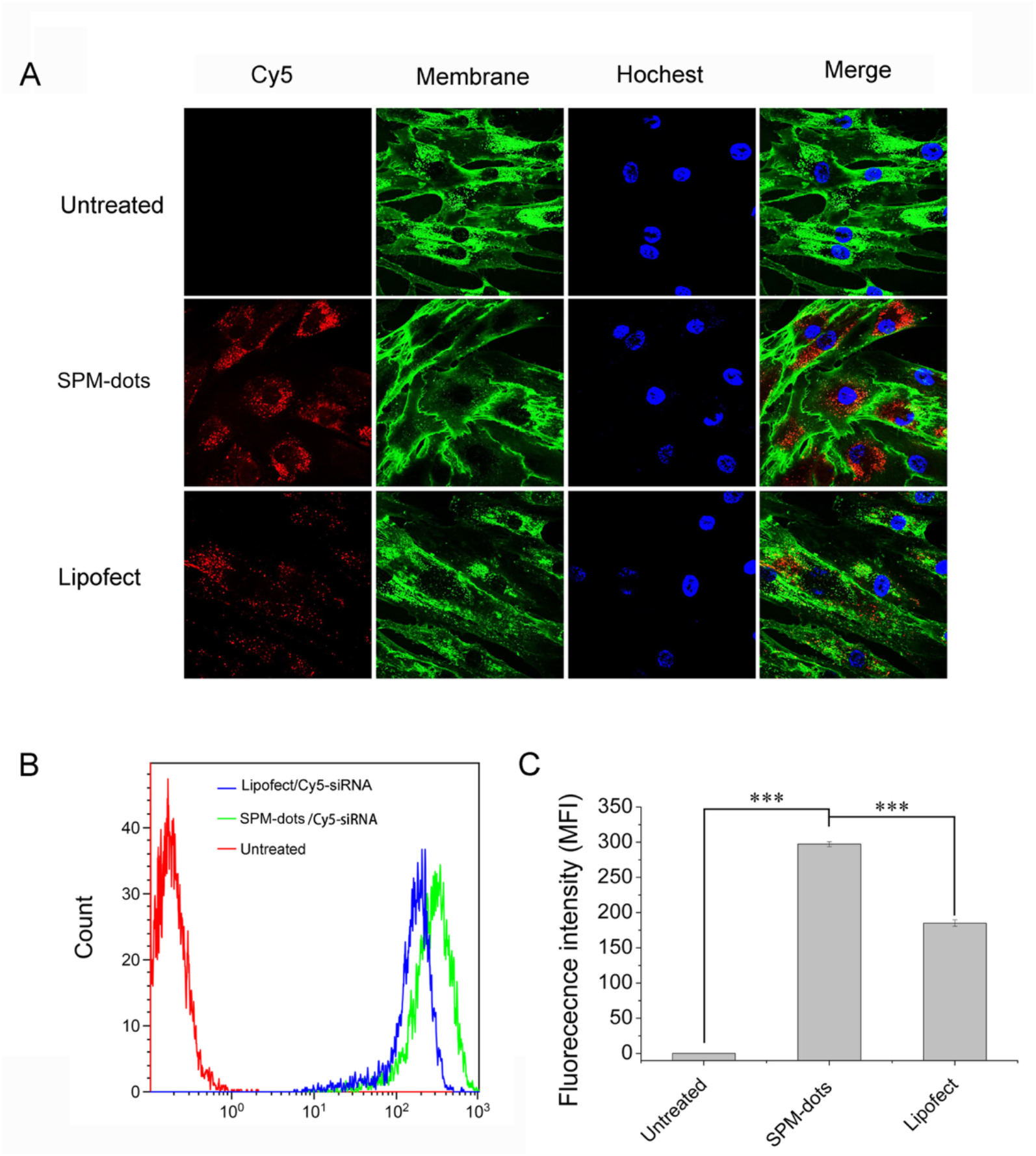
Intracellular uptake of SPM-dots/Cy5-siRNA by hMSCs. (A) Representative confocal images of cell transfected with Cy5-siRNA loaded SPM-dots or Lipofectamine 2000 (Lipofect), and compared with untreated cells. Cell membrane was stained with WGA 488 (green) and siRNA was labeled with Cy5 (red). (B) Representative histogram indicating the cellular uptake of Cy5-siRNA transfected by SPM-dots and (C) the quantification of Cy5-siRNA by flow cytometry. Data was expressed as mean ± SD (n=3).

### Gene silencing efficiency

Next, we tested the gene silencing potential of siRNAs delivered by SPM-dots using MSC cells stably expressing GFP. As shown in Figure 4A, treatment with SPM-dots/siGFP led to a stronger reduction in GFP fluorescence compared to the Lipofect/siGFP. The knockdown efficiency was also quantified by flow cytometry, showing >80% knockdown efficiency for SPM-dots versus approximately 50% for Lipofect (Figure 4B, C), thus validating the high transfection and gene silencing efficiency of SPM-dots.

**Figure 4.**
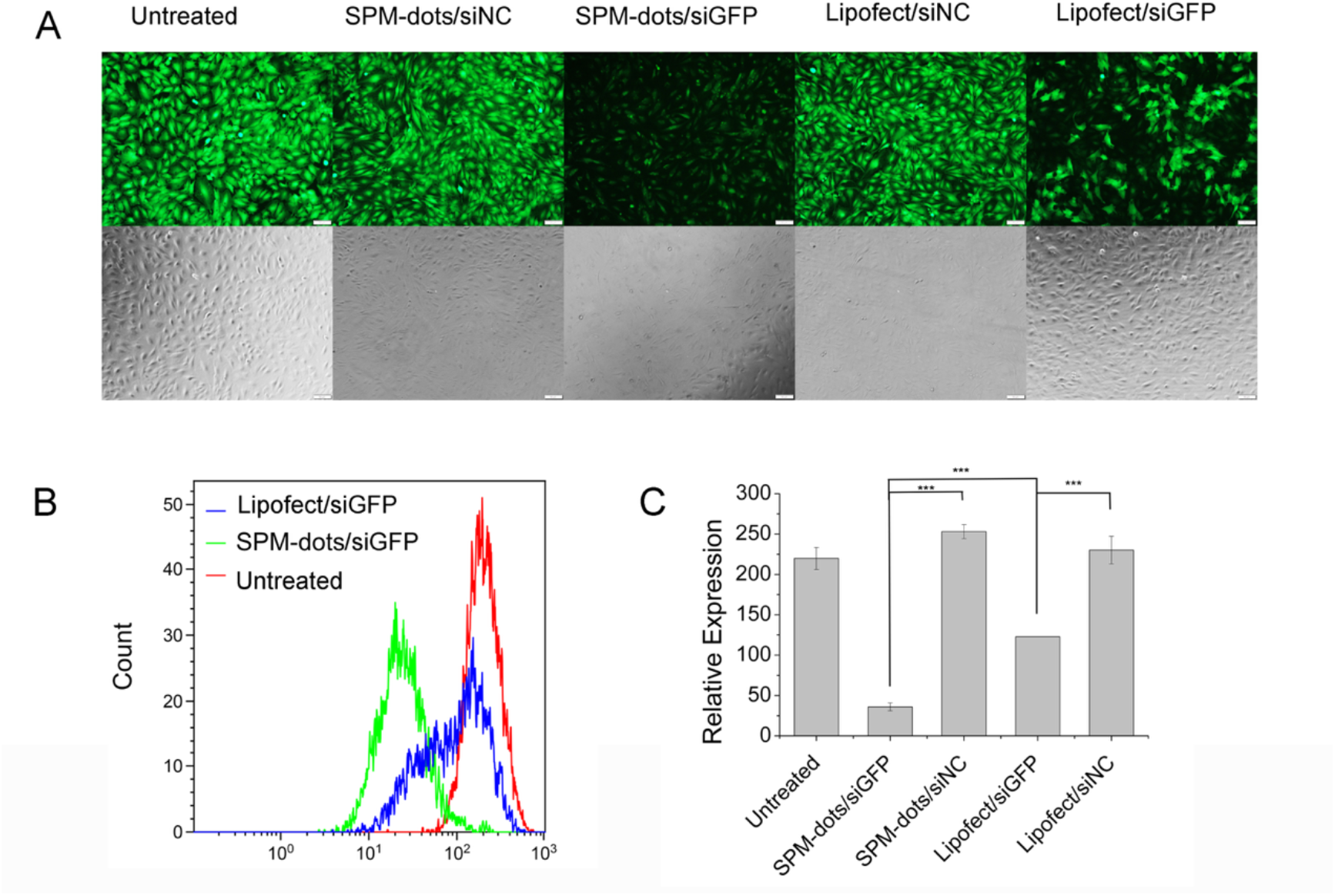
Silencing efficiency of SPM-dots/siRNA. MSCs stably expressing GFP were transfected with SPM-dots/siGFP or SPM-dots/siNC. Transfection efficiency was compared to commercial agent Lipofectamine 2000 (Lipofect). (A) Fluorescent images represent cells treated with indicated formulations after 48 hours. (B) Representative histogram indicating GFP florescence of MSCs after transfection. (C) GFP florescence intensity quantified by flow cytometry (mean ± SD, n=3) and normalized to untreated cells for comparison of silencing efficiency. *\*\*\*p<0*.*001*

### Effect of SPM-dots/miR-29b on osteoblast differentiation of hMSCs

MicroRNA-29b is known to be a key regulator of osteoblast differentiation^28^. To assess the effect of SPM-dots on osteoblast differentiation, primary hMSCs were transfected with SPM-dots/miR-29b or SPM-dots/miR-NC and subjected to osteogenic induction. After 14 days, *in vitro* mineralization was assessed by Alizarin Red S staining. As shown in Figure 5A, mineralization was markedly enhanced in cells treated with SPM-dots/miR-29b compared with controls. Quantitative analysis of mineral deposition was measured by relative absorbance, which showed approximately 2-fold enhanced mineralization in SPM-dots/miR29-b treated hMSCs relative to those treated with SPM-dots/miR-NC (Figure 5B).

**Figure 5.**
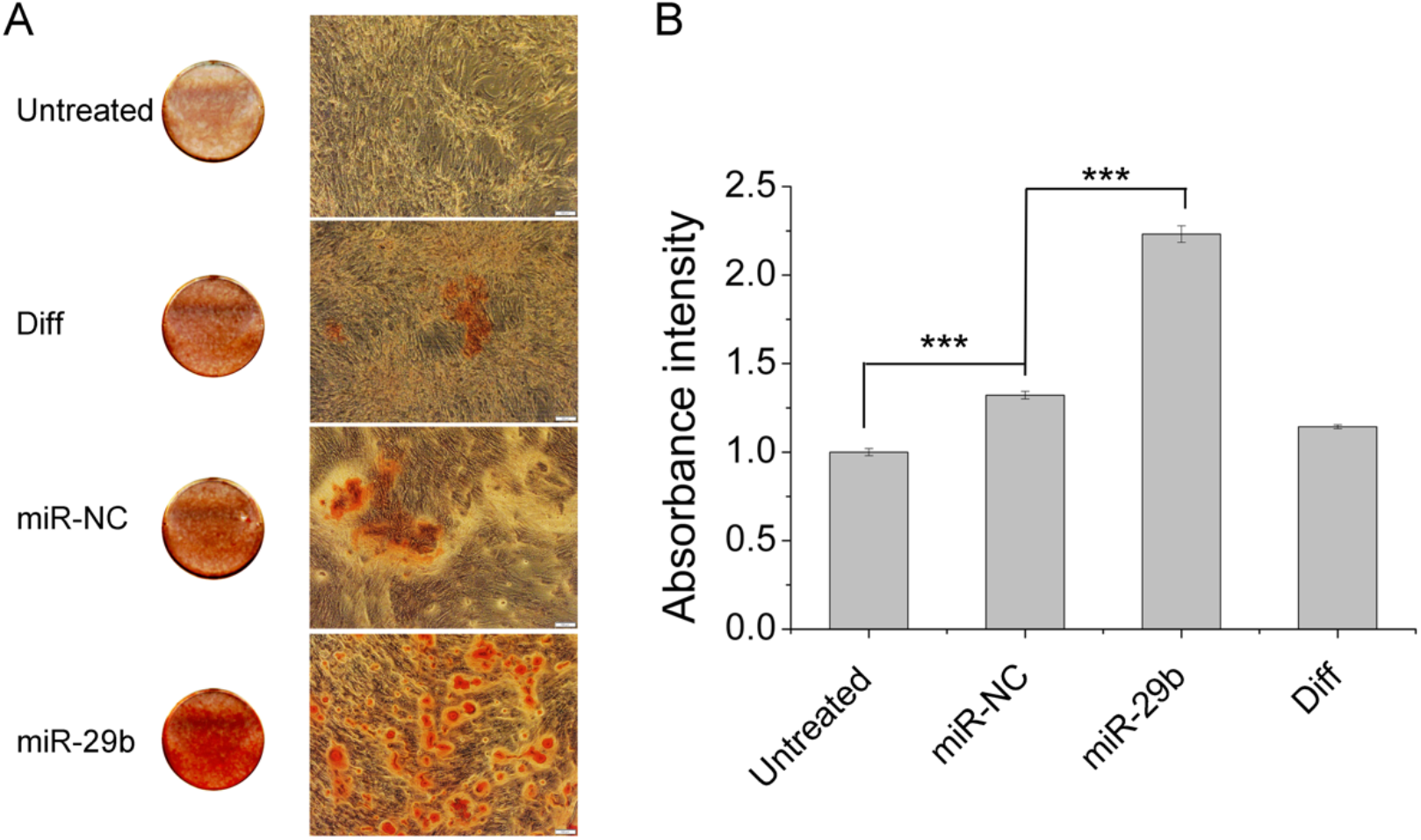
Effect of SPM-dots/miR-29b on osteogenic differentiation of hMSCs. hMSCs were transfected with SPM-dots/miR-29b or SPM-dots/miR-NC overnight and incubated in osteogenic media for 14 days. (A) Alizarin red staining of the untreated cells, differentiation medium alone, SPM-dots/miNC and SPM-dots/miR-29b in differentiation medium. (B) Quantification of in vitro mineralization was analyzed by the relative absorbance of eluted Alizarin Red S. Data was expressed as mean ± SD (n=3).

### Osteogenic-specific gene expression

To further validate the effect of the SPM-dots/miR-29b on osteoblast differentiation, expression levels of osteogenic-specific gene markers including RUNX2, OC, ALP and Col1A1 were quantified by q-PCR. Transfection with SPM-dots/miR-29b resulted in increased mRNA levels of RUNX2, OC and ALP, except for the down-regulation of Col1A1 (Figure 6). Notably, Col1A1 is one of the direct targets of miR-29b^29^, whereas the up-regulation of other markers reflects enhanced osteogenesis. As a late-stage osteogenic marker, OC expression was approximately 2-fold higher in SPM-dots/miR-29b treated cells compared with cells exposed to differentiation medium alone or SPM-dots/siNC.

**Figure 6.**
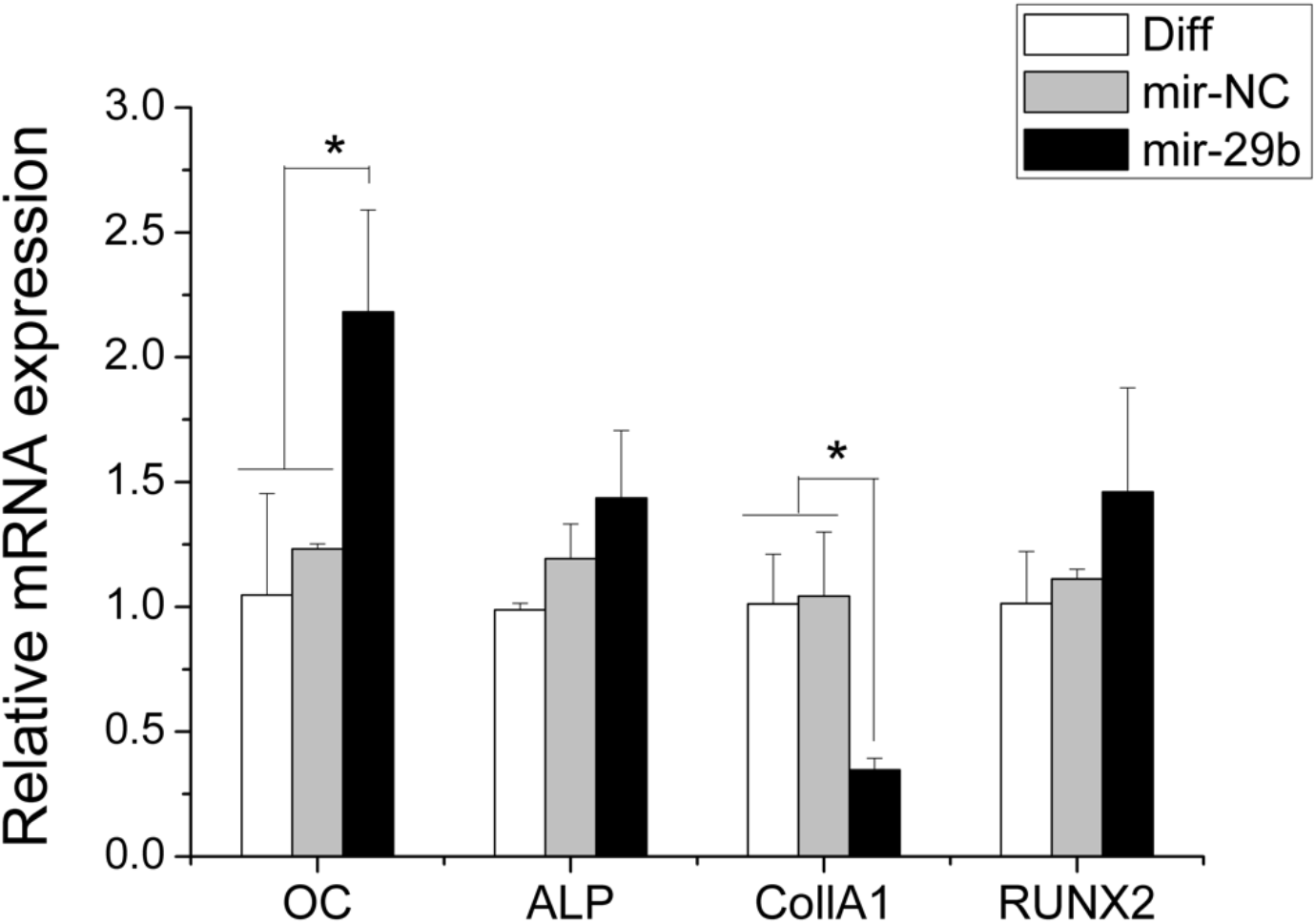
Osteogenic gene expression induced by SPM-dots/miR-29b. Relative expression of mRNA levels was analyzed by Real-time PCR and normalized to the group treated only with differentiation medium. Data was expressed as mean ± SD (n=3). * *p < 0*.*05*

### Stem cells tracking *in vivo*

To evaluate the cell tracking potential of SPM-dots, hMSCs transfected with SPM-dots/miRNA were mixed with Matrigel and implanted subcutaneously into a mouse. Live-animal imaging (Figure 7A) revealed strong fluorescent signals at the implantation sites, even with as few as 1×10^4^ implanted cells. Moreover, the relative total flux (photons/seconds) from the region of implantation correlated with the number of injected cells (Figure 7B).

**Figure 7.**
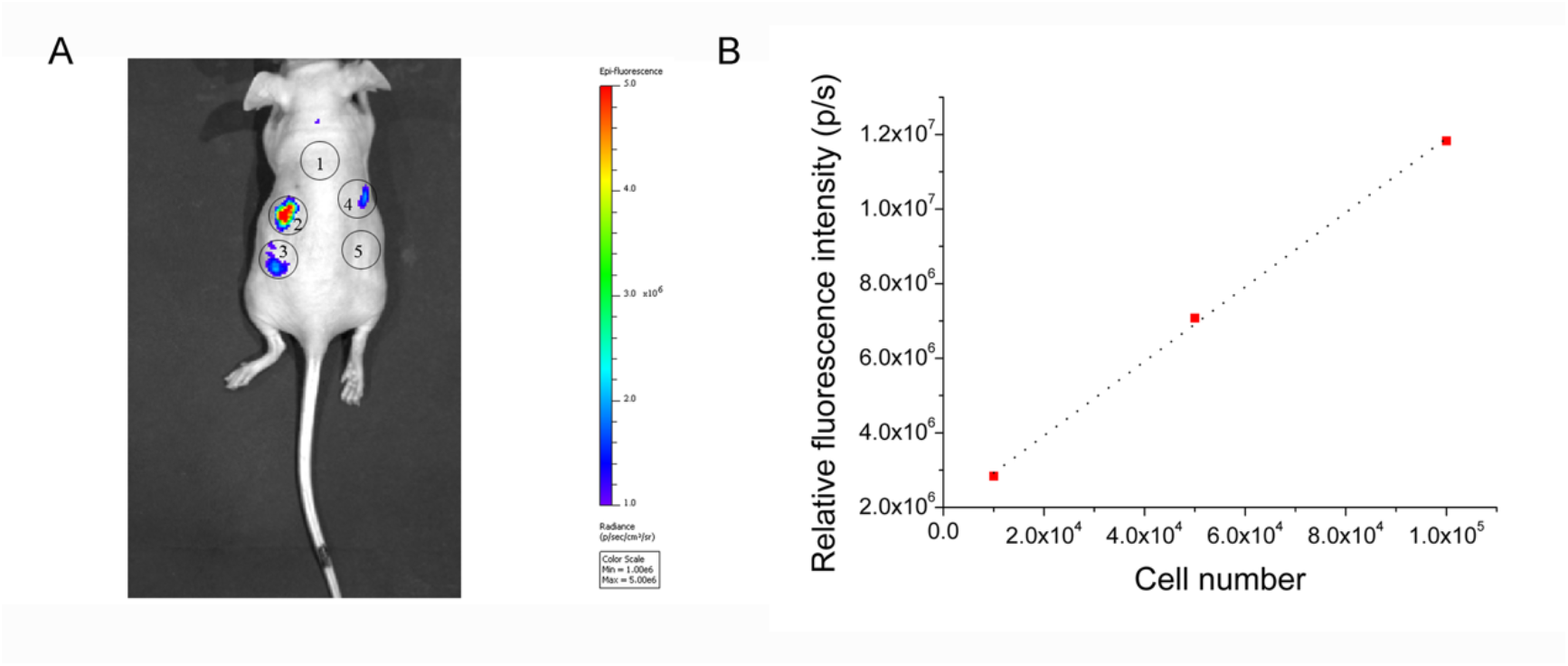
Stem cell labeling and *in vivo* tracking of hMSC cells labeled with SPM-dots /miRNA. (A) Different numbers of cells were mixed with Matrigel and the cells were subcutaneously implanted on the back of the Balb/c mouse. No.1 represents negative control (PBS/Matrigel), No.2: 1×10^5^, No.3 : 5×10^4^, No.4 : 1×10^4^ cells and No.5 : 5×10^3^ labeled cells. (B) Quantification of fluorescent signals in the regions of interest against cell number.

## Discussion

Various strategies have been used to promote MSC-dependent osteogenesis, including the delivery of growth factors, dexamethasone (Dex) and proteins such as bone morphogenetic proteins (BMPs)) ^30–33^. However, these approaches have notable limitations for the application, such as the Dex-induced inhibition of osteogenesis at high concentrations^34^, poor protein stability (BMPs), challenges in delivering large protein doses and the high cost^35^. In recent years, miRNAs have emerged as a promising alternative to induce osteogenesis. To enhance bone regeneration, researchers have engineered non-viral carriers such as chitosan nanoparticles^36^, and hydrogel^37^, peptide-based nanoparticles^38^ for delivery of miRNAs. Despite these advances, poor in vivo biocompatibility and low transfection efficiency remain significant challenges. The effectiveness of non-viral gene delivery in stem cells strongly depends on the specific type of vector used, which can significantly influence cellular uptake and osteogenic differentiation^39,40^. Additionally, biodistribution and migration of the injected MSCs are also essential for stem cell-based therapy. Several theranostic platforms that combine bioimaging and small RNA delivery have been reported in the literature, such as iron oxide based nanoparticles, quantum dots, gold nanoparticles, carbon nanotubes and nanoparticles loaded with dyes^41^. However, they are often associated with complex preparation protocols^42,43^, difficult detection^44^, toxicity^45^ or insufficient stability and low brightness^46,47^. Therefore, it is important to develop potent delivery systems that can overcome these barriers to combine miRNA delivery and stem cell tracking.

In this study, we have developed theranostic semiconducting polymer dots (SPM-dots) that integrate efficient gene delivery with robust cell-tracking capabilities. In our previous work, nanoparticles formulated with S14 and Pluronic F127 (FS14-NP) demonstrated excellent siRNA delivery efficacy in vitro and in vivo^16^. Pluronic F127 was selected due to its widespread use in pharmaceutical applications and approval by the U.S. Food and Drug Administration (FDA).

Building upon this system, we incorporated MEH-PPV into the system to create multifunctional SPM-dots with a condensed structure formed through hydrophobic interactions among S14, Pluronic F127 and MEH-PPV. In this design, S14 provides cationic groups for electrostatic interaction with RNA under acidic conditions, Pluronic F127 stabilizes the hydrophobic core and forms a hydrophilic corona for aqueous stability, and MEH-PPV serves as the fluorescence-emitting core of the system. MEH-PPV was chosen for its strong photoluminescence, biocompatibility and stability among conjugated polymers ^48–50^. Nanoparticles are typically considered stable when their zeta potential is over +30 mV or lower than −30 mV^51,52^. The SPM-dots exhibited a zeta potential of +41.2 mV with well-dispersed particle sizes, confirming their stability. Importantly, MEH-PPV incorporation did not compromise cell viability or miRNA encapsulation efficiency (Figure 2).

miR-29b is upregulated during osteogenic differentiation, leading to repression of various osteogenic inhibitors (such as, CTNNBIP1, histone deacetylase 4 (HDAC4), TGF-β3, and dual specificity phosphatase (DUSP2)) in order to promote osteogenesis and expression of osteogenic-specific markers such as ALP, RUNX2 and OC^53,54^. In our study, transfection with SPM-dots/miR29b enhanced in vitro matrix mineralization of hMSCs (Figure 5A, B). Although no statistically significant difference in ALP activity was observed among cells treated with SPM-dots/miR-29b, SPM-dots/mi-NC, or differentiation medium alone, ALP activity remained substantially higher than undifferentiated (untreated) cells, supporting ongoing osteogenic differentiation (data not shown; Figure S1). We also observed increased expression of osteogenic markers including RUNX2, OC and ALP. During the later stages of osteogenesis, miR-29b attenuates genes involved in mineralization regulation (e.g.COL1A1, COL5A3 and COL4A2 3’-UTR), facilitating extracellular matrix accumulation and organization^54^. Consistent with this, Col1A1 expression was downregulated by SPM-dots/miR-29b without negative impact on osteogenesis, as indicated by the enhanced mineralization. This suggests that miR-29b-mediated regulation of collagen expression may also avoid fibrosis during the late-stage mineralization. Finally, SPM-dots demonstrated high stem cell labeling efficiency and enabled in vivo tracking, with fluorescence intensity correlating with implanted cell numbers.

## Conclusion

In conclusion, we developed multi-functional semiconducting polymer dots (SPM-dots) that integrate miRNA/siRNA delivery with stem cell tracking. SPM-dots exhibited well-defined physicochemical properties and high gene silencing efficiency. Delivery of miR-29b significantly enhanced osteogenic differentiation, and SPM-dots enabled efficient stem cell labeling and in vivo tracking. This dual capability of targeted gene delivery and real-time cell monitoring highlights the potential of SPM-dots for applications in tissue regeneration and other therapeutic areas.

## Acknowledgement

This work is supported by Lundbeck Foundation (Project No. 23750) from Denmark. The authors also acknowledge the funding from RNA-Meta Center (NNF23OC0081177), the CEMBID program (NNF17OC0028070), and the China Scholarship Council (CSC) from China. The authors also thank Anne F. Nielsen for helpful comments on the manuscript.

