## supporting information for "Multifunctional semiconducting polymer dots as a miRNA delivery system for osteogenic differentiation and stem cell tracking"

**Table S1.** Primers used for mRNA amplification

| | Primers used for mRNA amplification |
|---|---|
| hGAPDH | F: 5'-GAAGGTGAAGGTCGGAGT-3'<br>R: 5'-GAAGATGGTGATGGGATTTC-3' |
| hALP | F: 5′-ACGTGGCTAAGAATGTCATC-3'<br>R: 5′-CTGGTAGGCGATGTCCTTA-3′ |
| hCOL1A1 | F: 5′-TGACGAGACCAAGAACTG-3′<br>R: 5′-CCATCCAAACCACTGAAACC-3′ |
| hOC (BGLAP) | F: 5′ CATGAGAGCCCTCACA 3′<br>R: 5′ AGAGCGACACCCTAGAC 3′ |
| hRUNX2 | F: 5'-CCATAACGGTCTTCACAAATCCT-3'<br>R: 5'-TCTGTCTGTGCCTTCTTGGTTC-3' |

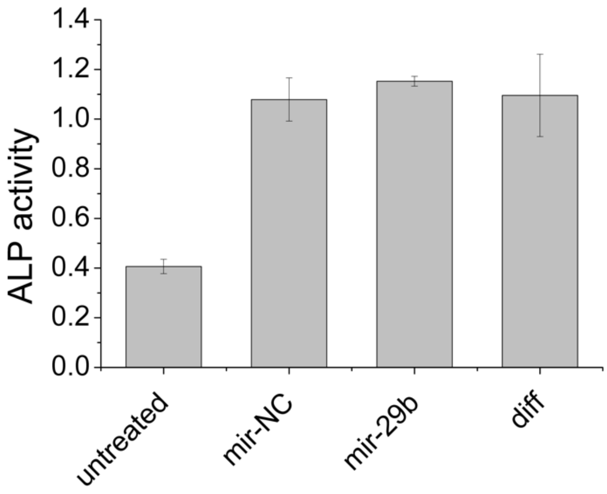


Figure S1. ALP activity of hMSCs treatment with SPM-dots/miR-29b, SPM-dots/miR-NC and compared with differentiation medium (diff) after osteoblast differentiation for 7 days.
